# Short-lived plants have stronger demographic responses to climate

**DOI:** 10.1101/2020.06.18.160135

**Authors:** Aldo Compagnoni, Sam Levin, Dylan Z. Childs, Stan Harpole, Maria Paniw, Gesa Römer, Jean H. Burns, Judy Che-Castaldo, Nadja Rüger, Georges Kunstler, Joanne M Bennett, C. Ruth Archer, Owen R. Jones, Roberto Salguero-Gómez, Tiffany M. Knight

**Affiliations:** Martin Luther University Halle-Wittenberg, Am Kirchtor 1, 06108, Halle (Saale), Germany; German Centre for Integrative Biodiversity Research (iDiv) Halle-Jena-Leipzig, Deutscher Platz 5e, Leipzig 04103, Germany; Department of Animal and Plant Sciences, University of Sheffield. Western Bank, Sheffield S10 2TN, UK; Department of Physiological Diversity, Helmholtz-Centre for Environmental Research –UFZ, Permoserstrasse 15, Leipzig 04318, Germany; Departamento de Biologia – IVAGRO, Universidad de Cadiz, Campus Rio San Pedro, 11510 Puerto Real, Spain; Interdisciplinary Centre on Population Dynamics (CPop), University of Southern Denmark. Campusvej 55, 5230 Odense M, Denmark; Department of Biology, University of Southern Denmark. Campusvej 55, 5230 Odense M, Denmark; Department of Biology, Case Western Reserve University, Cleveland, OH 44106-7080; Alexander Center for Applied Population Biology, Conservation & Science Department, Lincoln Park Zoo, Chicago, IL 60614-4712 USA; Smithsonian Tropical Research Institute, Apartado 0843-03092, Balboa, Ancón, Panama; Department of Economics, University of Leipzig, Grimmaische Straße 12, 04109 Leipzig, Germany; Univ. Grenoble Alpes, INRAE, UR LESSEM, 38000 Grenoble, France; Centre for Applied Water Science, Institute for Applied Ecology, The University of Canberra, Canberra, Australian Capital Territory, Australia; Centre for Ecology and Conservation, College of Life and Environmental Sciences, University of Exeter, Penryn, United Kingdom; Institute of Evolutionary Ecology and Conservation Genomics, University of Ulm, Albert-Einstein-Allee 11, 89081, Ulm, Germany; Department of Zoology, University of Oxford. 11a Mansfield Road, Oxford, OX1 3SZ, United Kingdom; Department of Community Ecology, Helmholtz Centre for Environmental Research–UFZ, 06120 Halle (Saale), Germany

**Keywords:** Climate change, COMPADRE Plant Matrix Database, Integral projection model (IPM), Life-history strategy, Matrix population model (MPM), PADRINO IPM Database, Precipitation, Temperature

## Abstract

To mitigate and adapt to climate change, there is an urgent need to synthesize the state of our knowledge on plant responses to climate. The availability of open-access data, combined with our understanding of plant physiology and life history theory provide opportunities to examine quantitative generalizations regarding which biomes and species are most responsive to climate drivers. Here, we synthesized time series of structured population models from 165 populations from 62 plants around the globe to link plant population growth rates to precipitation and temperature drivers. We expected: (1) more pronounced demographic responses to precipitation than temperature, especially in arid biomes; (2) a higher climate sensitivity in short-lived rather than long-lived species; and (3) a stronger response to climate by species that reproduce more frequently. We found that precipitation anomalies have a nearly three-fold larger effect on *λ* than temperature. Precipitation has substantially more pronounced effects in more arid sites, but large noise makes this relationship non-significant. Species with shorter generation time have much stronger absolute responses to climate anomalies, while the degree of iteroparity does not correlate with population responses to climate. We conclude that key species-level traits can predict plant population responses to climate, and discuss the relevance of this generalization for conservation planning and evolutionary theory.

## Introduction

Climate change is altering the mean as well as the variance in temperature and precipitation worldwide (IPCC 2013). These changes in climate are widely recognized as a prime global threat to biodiversity (Willis & Bhagwat 2009) because temperature and precipitation ultimately drive the demographic processes that determine the size and long-term viability of natural populations (Ehrlén & Morris 2015). Hence, it is critical to evaluate which species are most responsive to climatic drivers, and in which biomes (Sutherland *et al*. 2013). The urgency to understand the response of species to climate is particularly high for species that cannot buffer against the effects of climate change by migrating, such as sessile species. Among sessile species, numerous plants have short-distance dispersal, and cannot therefore shift their ranges fast enough to keep up with the current pace of climate change (Jump & Peñuelas 2005; Zhu *et al*. 2012).

Based on plant physiology, we expect that precipitation, or its interaction with temperature, predict plant population growth better than temperature alone. Most plant physiological processes, such as seed germination, tissue growth, floral induction, and seed set, are affected by water availability (Jones 2013). Temperature can also influence these processes, but typically by modulating water availability, as plant growth occurs across a wide range of temperatures (namely between 5° to 40° Celsius; Körner 2008, Jones 2013). The effect of temporal fluctuations on the growth rate of a population should be proportional to precipitation or temperature anomalies, where anomalies are deviations from mean values.

Precipitation and temperature anomalies are expected to have more pronounced effects in arid and cold biomes than in wet and temperate ones. Water availability, the difference between precipitation and evaporative demands, is known to be a strong predictor of vegetation types and structure (Stephenson 1990, Stegen et al. 2011). Accordingly, as water availability decreases, precipitation becomes the main factor limiting plant physiological processes (Noy-Meir 1973, Huxman et al. 2004). On the other hand, in cold biomes such as the tundra, temperature anomalies can dramatically change the length of the growing season (Bryson 1974).

Life-history theory also provides expectations for how natural plant populations may respond to climate drivers. Plant life-history strategies are defined by generation time, which describes how fast individuals in a population are substituted (and is correlated with life expectancy; Gaillard *et al*. 2005), and by the degree of iteroparity, which describes how many times individuals reproduce during their lifetimes (Salguero-Gómez *et al*. 2016; Salguero-Gómez 2017). The population growth of long-lived species should respond weakly to climatic anomalies compared to short-lived species. We expect this because the long-run population growth rate of long lived species respond less strongly to variation in survival, growth, and reproduction (Morris *et al*. 2008). There is less theory available to predict how the degree of iteroparity might influence the response of species to climatic anomalies (but see Demetrius 1978). However, frequent reproduction is expected to evolve as a bet-hedging strategy in unpredictable environments that allows populations to take advantage of years with favorable climate (Dewar & Richard 2007; Collett *et al*. 2018). Hence, we expect species with long reproductive windows (highly iteroparous species) to be more responsive to temporal climatic anomalies than species with short ones (semelparous species).

Understanding the effect of climate change on plant population growth rate requires demographic datasets that sample the climatic variability typically observed at a site (e.g. Tenhumberg *et al*. 2018). We expect that under-sampling this climatic variability will hinder the estimation of the effects of climate anomalies on population dynamics. However, it is unclear how long a demographic dataset must be to appropriately sample climatic variation that occurs at a population.

Here, we capitalize on the recent availability of large volumes of demographic data to quantitatively test how plant population growth rate, *λ*, responds to temperature and precipitation anomalies. We expect (*H*_*1*_) *λ* to be more strongly associated with precipitation than temperature anomalies, because we expect water availability to have stronger physiological effects than temperature; (*H*_*2*_) *λ* of plants in water-limited biomes to be more responsive to precipitation anomalies; (*H*_*3*_) *λ* of plants in cold biomes to be more responsive to temperature anomalies; (*H*_*4*_) species with greater generation time to respond more weakly to temperature and precipitation anomalies, (*H*_*5*_) species with a higher degree of iteroparity to respond more strongly to climatic anomalies. To contextualize our findings, we assessed how temporal replication affects our ability to sample climatic extremes.

## Methods

### Demographic data

To address our hypotheses, we used matrix population models (MPMs) or Integral projection models (IPMs) from the COMPADRE Plant Matrix Database (v. 5.0.1; Salguero-Gómez *et al*. 2015) and the PADRINO IPM Database (Levin et al. 2020). We selected models describing the transition of a population from one year to the next. Among these, we selected studies with at least six annual transition matrices, to balance the needs of adequate yearly temporal replicates and sufficient sample size of data for quantitative synthesis. We excluded density-dependent (8 studies) models, which we could not directly compare to the density-independent models (see next paragraph). However, one of these excluded studies provided raw data with high temporal replication (14 to 32 years of sampling) for 15 species (Chu *et al*. 2016). Therefore, we requested raw data from the authors of this study and re-analyzed these data to produce density-independent MPMs that were directly comparable to other studies (see methods for this in Appendix S1). The resulting dataset consisted of 46 studies, 62 species, 165 populations, and a total of 3,744 MPMs and 52 IPMs (Table S1). Almost all of these studies were conducted in North America and Europe (Fig. S1), in temperate biomes that are cold, dry or both cold and dry (Fig. S1, insert). The analyzed plant populations were tracked for a mean of 16 (median of 12) annual transitions.

We used the MPMs and IPMs in this dataset to calculate the response variable of our analyses: the yearly asymptotic population growth rate (*λ*). This measure is one of the most widely used summary statistics in population ecology (Sibly & Hone 2002), as it integrates the response of multiple interacting vital rates. Specifically, *λ* reflects the population growth rate that a population would attain if its vital rates remained constant through time (Caswell 2001). This metric therefore distills the effect of underlying vital rates on population dynamics, free of other confounding factors (e.g. transient dynamics arising from population structure; Stott et al. 2010). We calculated *λ* of each MPM or IPM with standard methods (Caswell 2001; Ellner *et al*. 2016). Because our MPMs and IPMs described the demography of a population transitioning from one year to the next, our *λ* values were comparable in time units. Finally, we identified and categorized any non-climatic driver associated with these MPMs and IPMs. Data associated with twenty one of our 62 species explicitly quantified a non-climatic driver (e.g., grazing, neighbor competition), for a total of 60 of our 165 populations. Of the datasets associated with these species, 19 included discrete drivers, and only three included a continuous driver.

### Climatic data

To test the effect of temporal climatic variation on demography, we gathered global climatic data. We downloaded 1 km^2^ gridded monthly values for maximum temperature, minimum temperature, and total precipitation between 1901 and 2016 from CHELSAcruts (Karger *et al*. 2017), which combines the CRU TS 4.01 (Harris *et al*. 2014) and CHELSA (Karger *et al*. 2017) datasets. These datasets include values from 1901 to 2016, which is necessary to cover the temporal extent of all 165 plant populations considered in our analysis. For our temperature analyses, we calculated mean monthly temperature as the mean of the minimum and maximum monthly temperatures. We used monthly values to calculate time series of mean annual temperature, and total annual precipitation at each site. We then used this dataset to calculate our “annual anomalies” for each census year, defined as the 12 months preceding a population census. Our annual anomalies are standardized z-scores. For example, if *X* is a vector of 40 yearly precipitation or temperature values, *E*() calculates the mean, and σ() calculates the standard deviation, we compute annual anomalies as *A* = [*X* - *E*(*X*)]/*σ*(*X*). Therefore, an anomaly of one refers to a year where precipitation or temperature was one standard deviation above the 40-year mean. In other words, anomalies represent how infrequent annual climatic conditions are at a site. Specifically, if we assume that A values are normally distributed, values exceeding one and two should occur every six and 44 years, respectively. We used 40-year means because the minimum number of years suggested to calculate climate averages is 30 (World Meteorological Organization 2017).

To test how the response of plant populations to climate changes based on biome we used two proxies of water and temperature limitation. For each study population, we quantified water limitation by computing a water availability index (WAI), and temperature limitation using mean annual temperature. To compute these metrics, we downloaded data at 1 km^2^ resolution for mean annual potential evapotranspiration, mean annual precipitation, and mean annual temperature referred to the 1970-2000 period. We obtained potential evapotranspiration data from the CGIAR-CSI consortium (http://www.cgiar-csi.org/). This dataset calculates potential evapotranspiration using the Hargreaves method (Zomer *et al*. 2008). We obtained mean annual precipitation and mean annual temperature from Worldclim (Hijmans *et al*. 2005). Here, we used WorldClim rather than CHELSA climatic data because the CGIAR-CSI potential evapotranspiration data was computed from the former. We calculated the WAI values at each of our sites by subtracting mean annual potential evapotranspiration from the mean annual precipitation (e.g. Vicente-Serrano *et al*. 2013).

### The overall effect of climate on plant population growth rate

To test *H*_*1*_, we estimated the overall effect sizes of response to anomalies in temperature, precipitation, and their interaction with a linear mixed effect model.

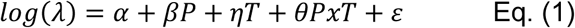

where *log(λ)* is the log of the asymptotic population growth rate of plant population *P is* precipitation, T is temperature. We included random population effects on the intercept and the slopes to account for the non-independence of measurements within populations. We then compared the mean absolute effect size of precipitation, temperature, and their interaction. This final model did not include a quadratic term of temperature and precipitation because these additional terms led to convergence issues. This likely occurred because single data sets did not include enough years of data.

### Population-level effect of climate on plant population growth rates

To test our remaining four hypotheses, we carried out meta-regressions where the response variable was the slope (henceforth “effect size”) of climatic anomalies on population growth rate for each of our populations. Before carrying out our meta-regressions, we first estimated the effect size of our two climatic anomalies on the population growth rate of each population separately. We initially fit population-level and meta-regressions simultaneously, in a hierarchical Bayesian framework. However, these Bayesian models shrunk the uncertainty of the most noisy population-level relationships, resulting in unrealistically strong meta-regressions. We therefore chose to fit population models separately, resulting in more conservative results. We fitted one of two types of model to each population:

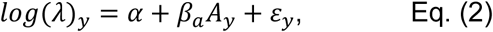

or

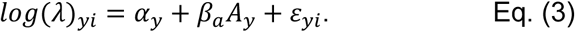

The model in equation 1 is a linear regression relating each *log(λ)* data point observed in year *y*, to the corresponding climate anomaly *A* observed in year *y*, via the intercept α, the effect size, β_a_, and a normally distributed error term, ε. The model in equation 3 is a mixed effect linear regression where *α*_*y*_ is a normally distributed random year effect. We fitted the mixed effect model in Eq. 3 only when multiple spatial replicates per each population were available each year. This happened in the few cases when a study contained contiguous populations, with no ecologically meaningful (e.g. habitat) differences. By using climate anomalies (*A*), we can interpret the effect sizes (*β*_*a*_) estimated in these models as a description of how plant populations respond to increasingly infrequent climatic conditions.

We included non-climatic covariates in our models whenever these substantially interacted with the climatic anomalies. A substantial interaction with a covariate would confound the effect of climate on *log(λ)*. To test for an interaction, we compared the corrected Akaike Information Criterion (AICc, Hurvich & Tsai 1989) between a model including one climatic anomaly (Eq. 2 or Eq. 3), and a model including a climatic anomaly, the covariate, and their interaction:

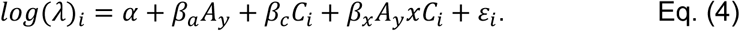

When AICc selected the model including the covariate (Eq. 4), we calculated the effect size of climate as *β*_*a*_ + *β*_*x*_*E*(*C*_*i*_). Hence, we added to *β*_*a*_ the effect of the interaction evaluated at the mean value of the covariate. For categorical variables comparing a control with a treatment, we calculate the effect size as *β*_*a*_ + *β*_*x*_0.5. We tested for an interaction between a covariate and climate anomaly in 18 of the 21 studies that included a non-climatic covariate. In the remaining three studies, covariates corresponded with the single populations. Because Eq. 2 and 3 are fit on separate populations, they implicitly accounted for covariates. In our subsequent analysis, only one population showed a substantial effect of the anomaly by covariate interactions (*Eryngium alpinum*, Table S1). We carried out these and subsequent analyses in R version 3.6.1 (R Core Team 2019).

### The effect of biome on the response of plants to climate

We used a simulation procedure to run two meta-regressions to test for the correlation of water and temperature limitation, and the effect size of climate drivers on *λ*. This meta-regression accounted for the uncertainty, measured as the standard error, in the effect sizes of climate drivers. We represented the effect of biome using a proxy of water (WAI) and temperature (mean annual temperature) limitation. The response data of this analysis were the effect sizes (*β*_*a*_ values) estimated by Eq. 2-4 for each of our 165 populations. In these meta-regressions the weight of each effect size was inversely proportional to its standard error. To test *H*_*2*_ and *H*_*3*_ on how biome should affect the response of populations to climate, we used linear meta-regressions. These two hypotheses tested both the sign and magnitude of the effect of climate. Therefore, we used the effect sizes as a response variable which could take negative or positive values. As predictors, we used population-specific WAI (*H*_*2*_, only for effect sizes quantifying the effect of precipitation), and mean annual temperature (*H*_*3*_, only for effect sizes quantifying the effect of temperature). Here, we note that because Eq. 2-4 estimated *β*_*a*_ values using A, which are *standardized* anomalies (z-scores), the meta-regressions testing H_2_ and H_3_ are affected by a small statistical artifact. This is especially true for the meta-regression testing *H*_*2*_. In this case, WAI is strongly correlated with the standard deviation of *absolute* (measured in mm) precipitation anomalies. If the per-mm effect of absolute precipitation anomalies did not change across sites, estimating the *β*_*a*_ values in Eq. 2-3 using z-scores will yield *β*_*a*_ values that increase, rather than decrease, with WAI. In Appendix S2 we describe this artifact, and we show that it has a negligible influence on our results.

We performed these two meta-regressions by exploiting the standard error of each effect size. We simulated 1,000 separate datasets where each effect size was independently drawn from a normal distribution whose mean was the estimated *β*_*a*_, and the standard deviation was the standard error of this *β*_*a*_. These simulated datasets accounted for the uncertainty in the *β*_*a*_ values. We fit 1,000 linear models, extracting for each its slope, *β*_*meta*_. Each one of these slopes had in turn its uncertainty, quantified by its standard error, *σ*_*meta*_. For each *β*_*meta*,_ we then drew 1000 values from a normal distribution with mean *β*_*meta*_ and standard deviation *σ*_*meta*_. We used the resulting 1 ×10^6^ values to estimate the confidence intervals of *β*_*meta*_. This procedure assumes that slopes are normally distributed. We performed a one-tailed hypothesis test, considering meta-regression slopes significant when over 95% of simulated values were below (*H*_*2*_) or above (*H*_*3*_) zero.

### The effect of life-history on the response of plants to climate

To test *H*_*4*_ and *H*_*5*_ on how the life-history of a species should mediate its responses to climate, we used gamma meta-regressions to model the absolute value of effect sizes (*β*_*a*_). Our response variable was |*β*_*a*_| because our predictions referred to the magnitude of responses to precipitation and temperature, separately. Using |*β*_*a*_| meant that, in these meta-regressions, the support of the response variable ranges from 0 to infinity. Thus, we fitted gamma meta-regressions using a log link. As predictors, we used generation time (*T*), and degree of iteroparity (*S*). We calculated *T* and *S* directly from the MPMs and IPMs using methods described in Appendix S3. We log-transformed both *T* and *S* to improve model fits. We carried out these meta-regressions using the same simulation procedure described for testing *H*_2_ and *H*_*3*._ We also carried out one-tailed hypothesis tests, by verifying whether 95% of *β*_*meta*_ values were below (*H*_*4*_) or above (*H*_*5*_) zero.

### Data quality

To quantify the relationship between study duration and the proportion of the climate sampled at a site, we first calculated the proportion of climatic variation sampled at a site in each of our populations. We define climate variation as the range of annual temperature and precipitation observed during the 40 years preceding the last census year of each dataset. At only one site (Chu et al. 2016), the last census year was 1935, so that this 40-year period extended prior to the first year of available climate data, which was 1901. At this site, we therefore characterized the climate variation using data for the 1901-1940 period. We then compared the range of climatic conditions experienced during each study period, to the range observed during the 40-year period representative of climate. We calculated this range of climatic conditions as the area of a two-dimensional convex hull enclosing the observed values of annual temperature and precipitation (Appendix S4). The proportion of climate sampled was the ratio between the range of climatic conditions observed during the demographic study of each population, and the range observed during the 40-year period representative of the climate of each population. We then quantified the relationship between the proportion of climate sampled, and the number of years sampled. We linked these two quantities by fitting a generalized additive model (Wood 2006) with three knots. This approach allows the line to saturate, while minimizing nonlinearities in the fitted line. We designed this analysis to quantify (i) the proportion of climate conditions sampled by studies at different durations and (ii) the minimum number of years it takes to reach the saturation point, and thus the optimal length of a study linking plant population growth rate to a climate driver.

## Results

### The overall effect of climate on plant population growth rate

As predicted (*H*_*1*_), the overall effect of precipitation anomalies on *log(λ)* was strong (β = 0.033, 95% C.I. [0.009,0.056]) relative to that of temperature (η = -0.012, 95% C.I. [- 0.034,0.010]) and their interaction (θ = -0.009, 95% C.I. [-0.029, 0.011]), which were centered around zero. On average, a year with precipitation one standard deviation above the mean changed *λ* by +3.3%.

### The effect of biome on the response of plants to climate

As expected (*H*_*2*_), populations responded more positively to precipitation anomalies in locations with lower water availability (Fig. 1A). On average, a year with precipitation one standard deviation above the mean would increase *λ* by roughly 6% in places with low water availability (WAI of -1500mm), and by 0% in places with higher water availability (WAI of 0mm). However, this relationship was not significant, with 92.8% of our bootstrap samples with slopes below zero (*β*_*meta*_ = -3.69 ×10^−5^, 95% C.I. [-8.67 ×10^−5^, 1.27 ×10^−5^]). We did not find evidence that the mean annual temperature (*H*_*3*_) of the site predicted the response of plant populations to temperature anomalies (Fig. 1B; *β*_*meta*_ = -1.30 ×10^−3^, 95% C.I. [-7.71 ×10^−3^, 5.08 ×10^−3^]).

**Figure 1.**
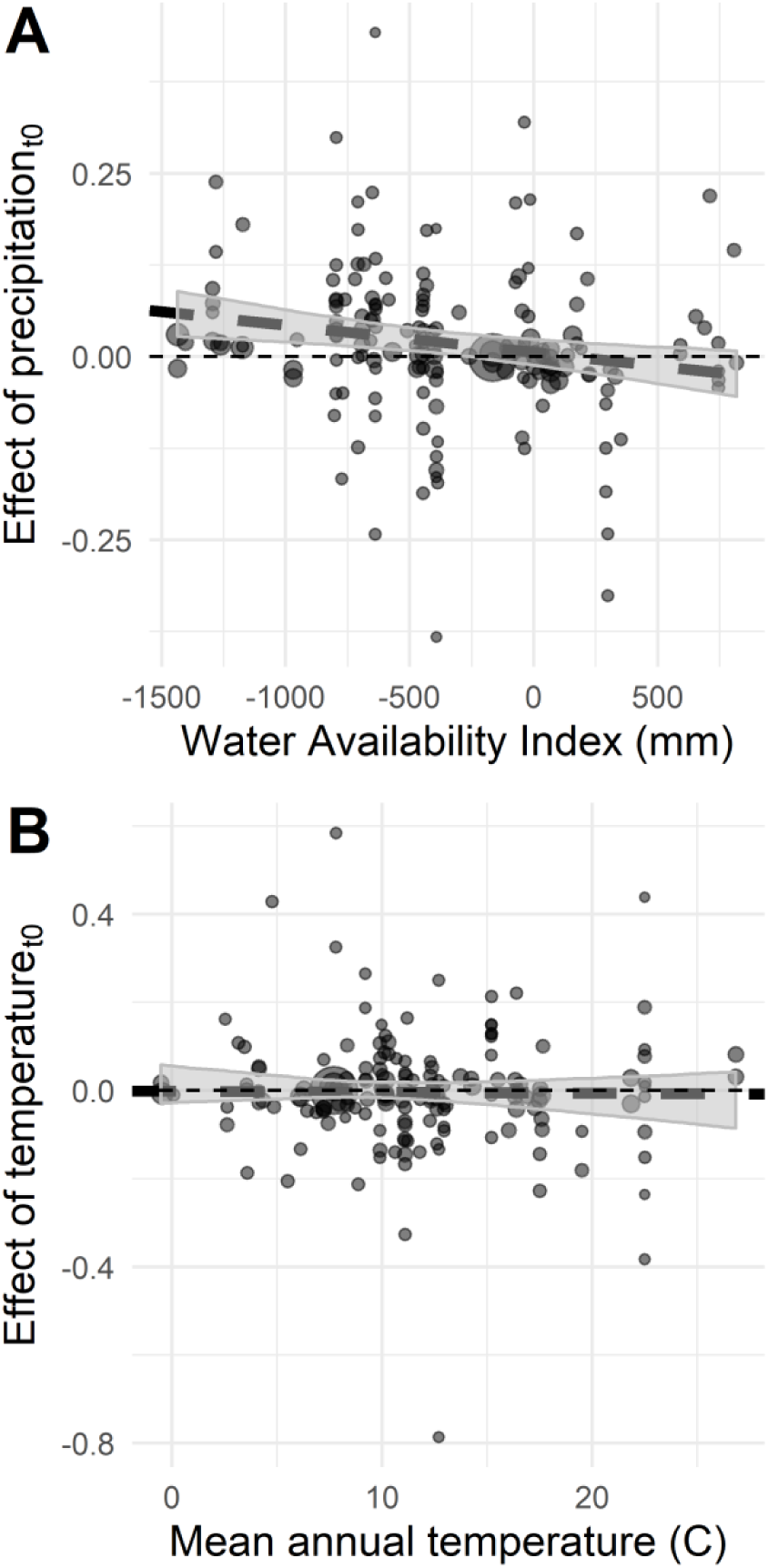
Effect of precipitation (**A**) and temperature (**B**) anomalies on the logged asymptotic population growth rate (*λ*) as a function of water availability index (**A**) and mean annual temperature (**B**). The y-axis represents the effect sizes of yearly anomalies in precipitation and temperature. The uncertainty of these effect sizes is shown by size of circles, which are inversely proportional to the standard error (SE) of effect sizes (1/SE). The shaded areas represent the 95% confidence interval of 1000 bootstrapped linear regressions. Solid lines show significant results, dashed lines show the zero effect.

### The effect of life-history strategy on the response of plants to climate

We found strong support for the effect of generation time (*H*_*3*_), and no support for the effect of the degree of iteroparity (*H*_*4*_) on the absolute response of plant populations to climate (Fig. 2). As expected, the response of species to climate correlated negatively with generation time (*H*_*4*_, Fig. 2A-B). In these meta-regressions, 100% of simulated *β*_*meta*_ values referring to the effect of precipitation (*β*_*meta*_ = -0.54, 95% C.I. [-0.71, -0.37]), and temperature (*β*_*meta*_ = -0.46, 95% C.I. [-0.64, -0.28]) were below zero. We found no relationship between the response of species to climate and the degree of iteroparity (Fig. 2C-D). In these two additional meta-regressions, the *β*_*meta*_ values associated with the effects of precipitation (*β*_*meta*_ = 0.001, 95% C.I. [-0.44, 0.40]) and temperature (*β*_*meta*_ = 0.09, 95% C.I. [-0.34, 0.52]) where above zero in only 50% and 67% of cases, respectively.

**Figure 2:**
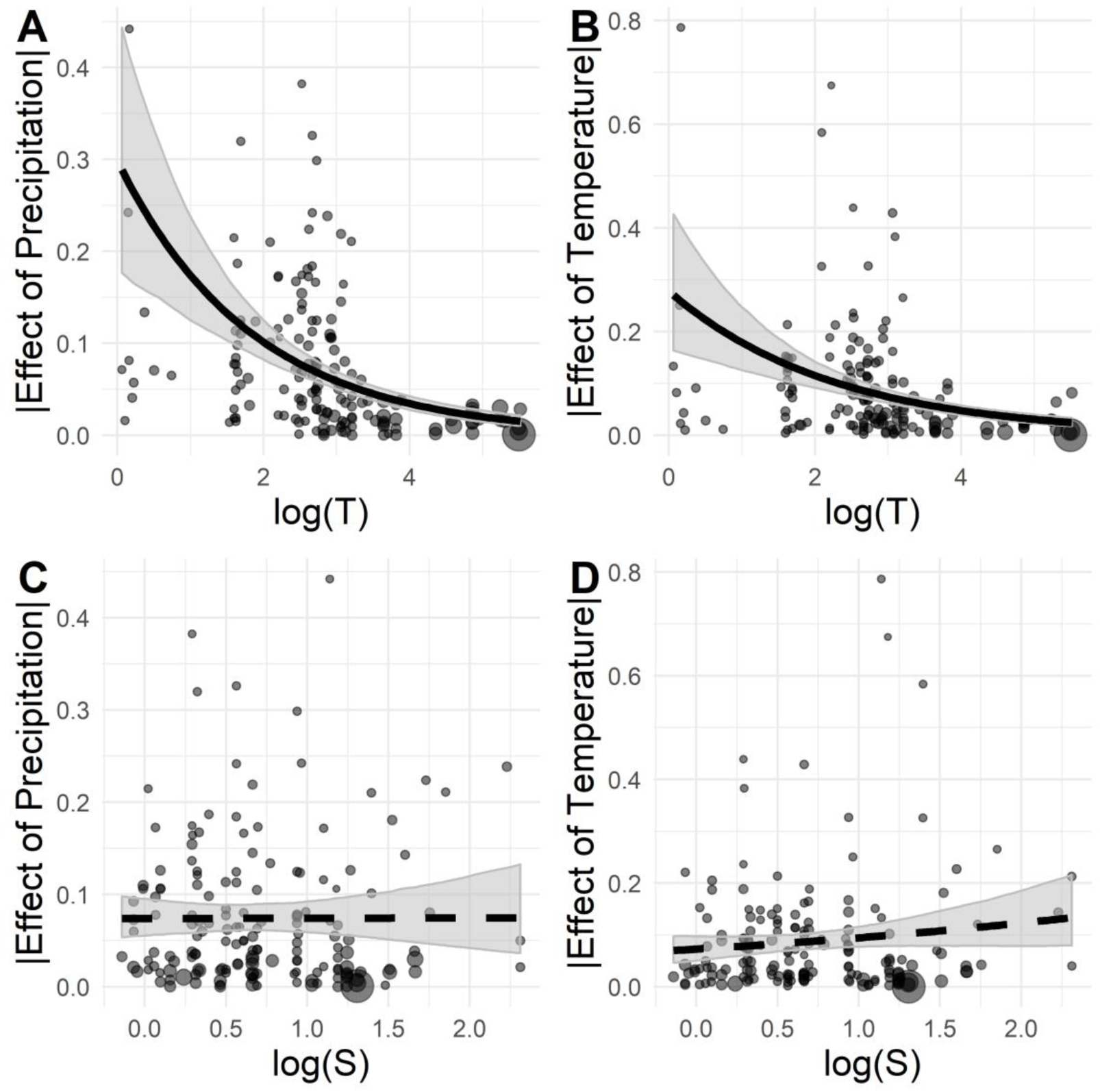
The absolute effect of precipitation and temperature as a function of logged generation time (*T*) and logged degree of iteroparity (*S*). We show the effect sizes of precipitation and temperature as a function of log(*T*) (**A** and **B**, respectively), and log(*S*) (**C** and **D**, respectively). The size of circles, shared areas, and solid versus dashed lines have the same meaning as in figure 1. However, in this case we fit gamma, rather than linear regressions.

### Proportion of climate sampled

The median and mean length of the existing long-term structured population studies available capture 41% and 58% of the climate variation that is likely to be experienced by plants at their study sites during a 40-year period. Six years of population censuses sampled less than 20% of the climatic conditions observed (Fig. 3). Our generalized additive model does not show a saturating asymptotic relationship between the number of MPMs (= sampled annual transitions), and the proportion of climate sampled by the study. However, the proportion of climate sampled decelerates after about 20 years (Fig. 3).

**Figure 3.**
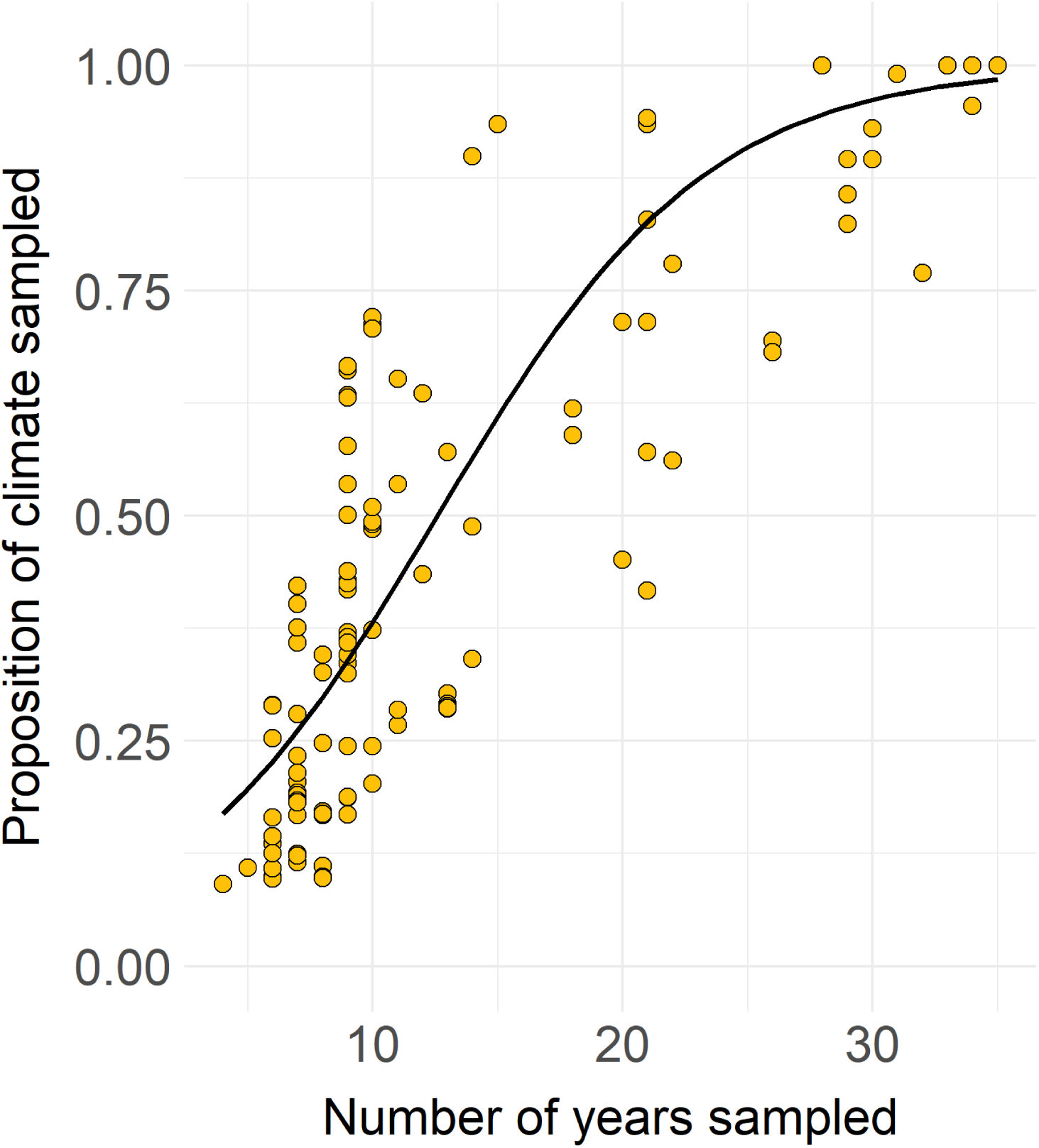
Fitted spline curve showing the relationship between the proportion of climate sampled and the numbers of years sampled by. Climate is defined as a two-dimensional area characterized by the extremes in annual mean temperature, and total precipitation observed during a 40-year period.

## Discussion

While understanding population responses to climate drivers has a long history in plant ecology (Andrewartha & Birch 1954), there is an urgent need to synthesize the state of our knowledge due to on-going climate change (Sutherland *et al*. 2013; IPCC 2014). The availability of open-access data (Hampton *et al*. 2013), a solid understanding of physiological ecology (Lambers *et al*. 2008), and a mature evolutionary theory of life histories (Hilde *et al*. 2020) provide opportunities to produce quantitative generalizations regarding plant population responses to climate. In our global synthesis, we found that (*H*_*1*_) precipitation has a stronger effect on population growth rates than temperature, and that (*H*_*4*_) plant species with shorter generation time respond more strongly to climate. Moreover, the magnitude of the non-significant relationship between water availability index and the response of plant populations to precipitation (*H*_*2*_) is ecologically large. We hypothesize this relationship is too noisy to be significant using an unstratified sample of 165 populations. Hence, the response of plant population dynamics to climate is more easily predicted at the species, rather than at the biome level, by using generation time. This novel generalization is relevant to conservation planning, and evolutionary theory.

Biome-level predictions of climate vulnerability remain elusive. The large, positive effect of precipitation on *log(λ)*, and the negative, smaller effects of temperature and its interaction with precipitation are consistent with the importance of water availability on plant population performance (Lambers *et al*. 2008). However, we also expected WAI to be a significant predictor of plant population responses to precipitation anomalies. Despite an ecologically large effect size, this relationship was non-significant (Fig. 1A). We expected responses to precipitation anomalies to correlate with WAI because the sensitivity to interannual variability in precipitation increases along aridity gradients for both annual net primary productivity (Huxman et al. 2004), and photosynthesis (Chen *et al*. 2019). While this relationship is likely to hold, it might be confounded by several factors. For example, indirect biotic effects mediated by precipitation could cause unexpected results (Gilman *et al*. 2010; Alexander *et al*. 2015; Rysavy *et al*. 2016). Moreover, the position of each species within its range will play a substantial role. Species located at their range edges should have a stronger response to climatic fluctuations (Kleinhesselink & Adler 2018; Fredston-Hermann *et al*. 2020). Finally, factors like longevity (see below) might de-couple the forcing effect of precipitation on population growth.

Our finding that the effects of temperature anomalies on plant population growth are weak and do not change based on mean annual temperature supports our hypothesis on the importance of water limitation, but it might also reflect that most of our populations were located in temperate biomes. In temperate systems, interannual fluctuations in temperature anomalies are not the main limiting factor for plant productivity (Schultz & Halpert 1993; Wang *et al*. 2003). The bias in existing studies towards temperate biomes might also explain why our models did not detect a strong interaction between temperature and precipitation. Many plant adaptations to high temperatures have evolved to limit water loss (Lambers *et al*. 2008; Jones 2013). Therefore, we might expect a strong interaction between precipitation and temperature anomalies where mean precipitation is low and mean temperature high. These conditions should occur, for example, in subtropical desert or tropical savannas, but only a handful of our studies occur in these biomes (Fig. S1).

Climate change projections involving precipitation are much more uncertain than those involving temperature (IPCC 2014). Moreover, prediction uncertainty in climate projection is not expected to improve much in the coming decades (Knutti & Sedláček 2013). Given that precipitation is the strongest predictor of plant population responses in our analyses, this implies that ecological forecasts will suffer from high uncertainty. As a result, ecological forecasts on plant populations might have to put great care in accounting for the uncertainty derived from climate change projections (e.g. “model uncertainty”, Dietze 2017).

Our results are the first to show that generation time is linked to population responses to climatic drivers across a large number of species. To our knowledge, the only other study to test for this hypothesis found a similar pattern for three amphibian species (Cayuela *et al*. 2017). The hypothesis linking generation time to population responses to climate was initially inferred from patterns of variation in the vital rates and population growth of plant and animal populations (Morris et al. 2008). Long generation time occurs in species that invest more in survival than in development and reproduction (Franco & Silvertown 2004). The negative relationship we found between generation time and climatic effects suggests that investment in survival could be an adaptation to buffer plant populations growth against climatic fluctuations (Fridley 2017). Hence, we demonstrated that it is possible to use plant traits to predict which species will be most sensitive to climate change (Sutherland *et al*. 2013). Interestingly, generation time is a fundamental quantity in identifying extinction probability (IUCN 2019; Staerk *et al*. 2019). It is therefore good news that this trait can also predict the climatic sensitivity of plants.

We speculate that the non-existing relationship between the degree of iteroparity and population responses to climate might reflect that reproduction responds to subtle climatic signals. We originally hypothesized that the degree of iteroparity is a bet-hedging strategy allowing species to take advantage of conditions favorable to reproduction. However, the reproduction of plants does not necessarily respond well to annual climate anomalies. For example, seed germination might be driven by specific environmental cues (Baskin & Baskin 1998; Levine *et al*. 2008). Similarly, tree masting responds to climate with a two-year lag (Moreira *et al*. 2015; Vacchiano *et al*. 2017). However, no study, to our knowledge, has systematically compared the inherent predictability of separate vital rates.

Improving our estimates of climate effects on *λ* might be best accomplished by increasing the spatial replication of demographic censuses. Our finding that even 12 year-long studies sample only 50% of climatic anomalies at a site, indicate that including multiple spatial, rather than temporal replicates might be a more strategic sampling strategy. Increasing the temporal replication of studies is efficient for populations that have been censused for a few years already. For example, adding five years of sampling to a pre-existing ten-year dataset would increase the coverage of climatic anomalies observed during a 40-year period from 37% to 62%. For studies established anew, investigators could sample multiple populations located relatively far from each other, thereby maximizing the coverage of species climatic spaces. Note that here we are not advocating inferring a temporal process from a spatial pattern, as in classic space for time substitutions (e.g. Blois *et al*. 2013). Rather, more spatial replicates should just increase our power to estimate the effect of climate. One final hurdle in the estimation of climate effects is that microclimatic conditions might substantially change how plants experience climate (Scherrer & Körner 2010). Hence, a way to improve estimates of climate effects is measuring, or estimating, how the climate above the ground relates to the microclimate experienced by plants (e.g. Souther & McGraw 2014; Körner *et al*. 2016).

Ecologists are invested with the challenge of forecasting how species abundance will change as a result of climate change. This need for prediction has led to the considerable popularity of species distribution models (Elith & Leathwick 2009). These models predict the geographic distribution of species by correlating species abundance data with environmental factors (e.g. average climatic conditions), but without understanding the underlying mechanisms. For example, (Csergő *et al*. 2017) showed that predicting species presence/absence from species distribution models and from MPMs results in a great deal of disagreement. These authors suggest that understanding how the environment affects the underlying demography of plants is a much more promising way to understand which species will persist in the future. In this study, we have risen to this challenge, providing the first generalizations on the climate sensitivity of plant populations.

## Acknowledgements

This article is the result of working group sAPROPOS (Analyses of PROjectionS of POpulations) supported by sDiv, the Synthesis Centre of the German Centre for Integrative Biodiversity Research (iDiv) Halle-Jena-Leipzig (funded by the German Research Foundation, FZT 118 - 202548816), led by R.S.-G. & T.M.K. T.M.K., A.C., and S.L. were supported by the Alexander von Humboldt foundation; R.S.-G. was supported by NERC IRF (NE/M018458/1); JC-C, RS-G, and ORJ were also supported by an NSF Advances in Biological Informatics grant (DBI-1661342000), N.R. was funded by a research grant from Deutsche Forschungsgemeinschaft DFG (RU 1536/3-1). We acknowledge the efforts of the Max Planck Institute for Demographic Research in curating and making the COMPADRE Plant Matrix Database open-access, as well as the numerous authors who have kindly shared their demographic data and population models.

## Supporting information

**Table S1.** List of studies, and associated plant species, providing the Matrix Population Models (MPMs) or Integral Projection Models (IPMs) used in this manuscript. We provide information on the originating article, the spatio-temporal replication of the study, the inclusion in the COMPADRE and PADRINO databases, and the type of non-climatic covariates linked to each population model. We used these archived projection models in our comparative analyses (Fig. 1 and 2).

| Species | Years | Subpop | Database | Article | Type of non-climatic covariate (if any) |
| --- | --- | --- | --- | --- | --- |
| *Actaea spicata* | 6 | 2 | COMPADRE | Fröborg; Eriksson 2003 Can J Bot |  |
| *Arabis fecunda* | 6 | 1 | COMPADRE | Lesica; Shelly 1995 Am J Bot |  |
| *Artemisia tripartita* | 22 | 6 |  | Chu et al. 2016 Nat Comm | mowing/grazing |
| *Asplenium scolopendrium* | 21 | 3 | COMPADRE | Bremer; Jongejans 2010 Popul Ecol |  |
| *Astragalus cremnophylax* var. *cremnophylax* | 6 | 1 | COMPADRE | Maschinski; Frye; Rutman 1997 Cons Biol | trampling |
| *Astragalus scaphoides* | 11 | 3 | COMPADRE | Crone; Lesica 2004 Ecology |  |
| *Astragalus scaphoides* | 26 | 4 | COMPADRE | Tenhumberg; Crone; Ramula; Tyre 2018 Ecol |  |
| *Astragalus tyghensis* | 9 | 5 | COMPADRE | Kaye; Pyke 2003 Ecology |  |
| *Bencomia exstipulata* | 9 | 1 | COMPADRE | Marrero; Oostermeijer;<br>Nogales; Van Hengstum;<br>Saro; Carqué; Sosa; Bañares<br>2019 J Nat Conserv |  |
| *Bouteloua curtipendula* | 34 | 2 | COMPADRE | Chu et al. 2016 Nat Comm |  |
| *Bouteloua eriopoda* | 18 | 7 | COMPADRE | Chu et al. 2016 Nat Comm |  |
| *Bouteloua eriopoda* | 32 | 10 | COMPADRE | Chu et al. 2016 Nat Comm |  |
| *Bouteloua gracilis* | 13 | 6 | COMPADRE | Chu et al. 2016 Nat Comm |  |
| *Bouteloua hirsuta* | 34 | 2 | COMPADRE | Chu et al. 2016 Nat Comm |  |
| *Bouteloua rothrockii* | 18 | 7 | COMPADRE | Chu et al. 2016 Nat Comm |  |
| *Brassica insularis* | 9 | 4 | COMPADRE | Noel; Maurice; Mignot;<br>Glémin; Carbonell; Justy;<br>Guyot; Olivier; Petit 2010<br>Cons Genet |  |
| *Centaurea corymbosa* | 6 | 3 | COMPADRE | Fréville; Colas; Riba; Caswell;<br>Mignot; Imbert; Olivier 2004<br>Ecology |  |
| *Centaurea corymbosa* | 21 | 6 | COMPADRE | Belaid; Maurice; Fréville;<br>Carbonell; Imbert 2018 Bio<br>Con |  |
| *Cirsium pitcheri* | 10 | 3 | COMPADRE | Ellis; Williams; Lesica; Bell;<br>Bierzychudek; Bowles; Crone;<br>Doak; Ehrlén; Ellis-Adam;<br>McEachern; Ganesan;<br>Latham; Luijten; Kaye; Knight;<br>Menges; Morris; Den Nijs;<br>Oostermeijer; Quintana-Ascencio; Shelly; Stanley;<br>Thorpe; Ticktin; Valverde;<br>Weekley 2012 Ecology |  |
| *Cirsium pitcheri* | 7 | 1 | COMPADRE | Bell; Powell; Bowles 2013 J Wild Manag |  |
| *Cirsium pitcheri* | 14 | 1 | COMPADRE | Jolls; Marik; Hamze; Havens 2015 Biol Cons |  |
| *Cirsium pitcheri* | 21 | 5 | COMPADRE | Halsey; Bell; McEachern; Pavlovic 2016 Ecosphere |  |
| *Cirsium undulatum* | 30 | 1 | COMPADRE | Dalgleish; Koons; Adler 2010 J Ecol |  |
| *Cryptantha flava* | 14 | 1 | COMPADRE | Salguero-Gómez; Kempenich; Forseth; Casper 2014 Ecology | climate |
| *Cypripedium calceolus* | 8 | 3 | COMPADRE | Nicolè; Brzosko; Till-Bottraud 2005 J Ecol |  |
| *Cypripedium fasciculatum* | 8 | 3 | COMPADRE | Thorpe; Stanley; Kayne; Latham 2011 Report |  |
| *Daphne rodriguezii* | 9 | 4 | COMPADRE | Rodriguez-Perez; Traveset 2012 Oikos | mutualists |
| *Dicerandra frutescens* | 12 | 7 | COMPADRE | Menges; Quintana-Ascencio; Weekley; Gaoue 2006 Biol Cons | disturbance |
| *Dracocephalum austriacum* | 7 | 6 | PADRINO | Nicolè et al. 2011 J Ecol |  |
| *Echinacea angustifolia* | 31 | 1 | COMPADRE | Dalgleish; Koons; Adler 2010 J Ecol |  |
| *Eriogonum longifolium* var. *gnaphalifolium* | 7 | 1 | COMPADRE | Satterthwaite; Menges; Quintana-Ascencio 2002 Ecol Appl | disturbance |
| *Eryngium alpinum* | 9 | 7 | COMPADRE | Andrello; Bizoux; Barbet-Massin; Gaudeul; Nicolè; Till-Bottraud 2012 Biol Cons | mowing/grazing |
| *Eryngium alpinum* | 9 | 2 | COMPADRE | Andrello; Devaux; Quétier;<br>Till-Bottraud 2018 J Appl Ecol | mowing/grazing |
| *Eryngium cuneifolium* | 9 | 3 | COMPADRE | Menges; Quintana-Ascencio<br>2004 Ecol Monog | disturbance |
| *Frasera speciosa* | 34 | 1 | COMPADRE | Che-Castaldo; Inouye 2011<br>Ecosphere |  |
| *Haplopappus radiatus* | 9 | 10 | COMPADRE | Kaye; Pyke 2003 Ecology |  |
| *Hedyotis nigricans* | 29 | 1 | COMPADRE | Dalgleish; Koons; Adler 2010 J<br>Ecol |  |
| *Helianthella quinquenervis* | 14 | 1 | COMPADRE | Compagnoni et al. 2016, Eco<br>Monog |  |
| *Helianthemum juliae* | 9 | 1 | COMPADRE | Marrero-Gómez;<br>Oostermeijer; Carqué-Álamo;<br>Bañares-Baudet 2007 Biol<br>Cons |  |
| *Hesperostipa comata* | 22 | 6 |  | Chu et al. 2016 Nat Comm | mowing/grazing |
| *Hesperostipa comata* | 13 | 6 |  | Chu et al. 2016 Nat Comm |  |
| *Horkelia congesta* | 6 | 1 | COMPADRE | Kaye; Benfield 2004 Report |  |
| *Kosteletzkya pentacarpos* | 8 | 1 | COMPADRE | Pino; Picó; Roa 2007 Bot J Lin<br>Soc |  |
| *Lesquerella ovalifolia* | 28 | 1 | COMPADRE | Dalgleish; Koons; Adler 2010 J<br>Ecol |  |
| *Lomatium bradshawii* | 7 | 7 | COMPADRE | Kaye; Pyke 2003 Ecology |  |
| *Opuntia imbricata* | 10 | 1 | PADRINO | Compagnoni et al. 2016, Eco<br>Monog |  |
| *Opuntia macrorhiza2* | 6 | 2 | COMPADRE | Haridas; Keeler; Tenhumberg<br>2015 Ecology |  |
| *Opuntia rastrera* | 7 | 2 | COMPADRE | Mandujano; Montaña;<br>Franco; Golubov; Flores-Martinez 2001 Ecology | site condition |
| *Orchis purpurea* | 6 | 6 | COMPADRE | Jacquemyns; Brys; Jongejans 2010 Ecology | site condition |
| *Panax quinquefolius* | 9 | 2 | COMPADRE | Chandler; McGraw 2017 J Ecol |  |
| *Paronychia jamesii* | 33 | 1 | COMPADRE | Dalgleish; Koons; Adler 2010 J Ecol |  |
| *Pascopyrum smithii* | 13 | 6 |  | Chu et al. 2016 Nat Comm |  |
| *Pediocactusbradyi* | 22 | 4 | COMPADRE | Shryock; Esque; Hughes 2014 Am J Bot |  |
| *Phyllanthus emblica* | 10 | 1 | COMPADRE | Ticktin; Ganesan; Paramesha; Setty 2012 J Appl Ecol | neighbors harvesting |
| *Phyllanthus indofischeri* | 10 | 1 | COMPADRE | Ticktin; Ganesan; Paramesha; Setty 2012 J Appl Ecol | neighbors harvesting |
| *Poa secunda* | 22 | 6 |  | Chu et al. 2016 Nat Comm | mowing/grazing |
| *Poa secunda* | 13 | 6 |  | Chu et al. 2016 Nat Comm |  |
| *Primula farinosa* | 5 | 2 | COMPADRE | Toräng; Ehrlén; Ågren 2010 Oecologia | site condition |
| *Primula veris* | 9 | 2 | COMPADRE | Jacquemyn; Brys; Davison; Tuljapurkar; Jongejans 2011 Oikos | mowing/grazing |
| *Pseudoroegneria spicata* | 22 | 6 |  | Chu et al. 2016 Nat Comm | mowing/grazing |
| *Psoralea tenuiflora* | 33 | 1 | COMPADRE | Dalgleish; Koons; Adler 2010 J Ecol |  |
| *Purshia subintegra* | 7 | 2 | COMPADRE | Maschinski; Baggs; Quintana-Ascencio; Menges 2006 Cons Biol | site condition |
| *Pyrocoma radiata* | 10 | 4 | COMPADRE | Pfingsten 2013 PhD thesis | site condition |
| *Ratibida columnifera* | 29 | 1 | COMPADRE | Dalgleish; Koons; Adler 2010 J Ecol |  |
| *Schizachyrium scoparium* | 34 | 2 |  | Chu et al. 2016 Nat Comm |  |
| *Serapias cordigera* | 13 | 6 | COMPADRE | Pellegrino; Bellusci 2014 Bot J Lin Soc | anthropic |
| *Silene spaldingii* | 12 | 1 | COMPADRE | Lesica; Crone 2007 J Ecol |  |
| *Solidago mollis* | 30 | 1 | COMPADRE | Dalgleish; Koons; Adler 2010 J Ecol |  |
| *Sphaeralcea coccinea* | 35 | 1 | COMPADRE | Dalgleish; Koons; Adler 2010 J Ecol |  |
| *Sporobolus flexuosus* | 32 | 10 |  | Chu et al. 2016 Nat Comm |  |
| *Thelesperma megapotamicum* | 29 | 1 | COMPADRE | Dalgleish; Koons; Adler 2010 J Ecol |  |
| *Trillium ovatum* | 6 | 2 | COMPADRE | Ream 2011 PhD thesis |  |

**Figure S1.**
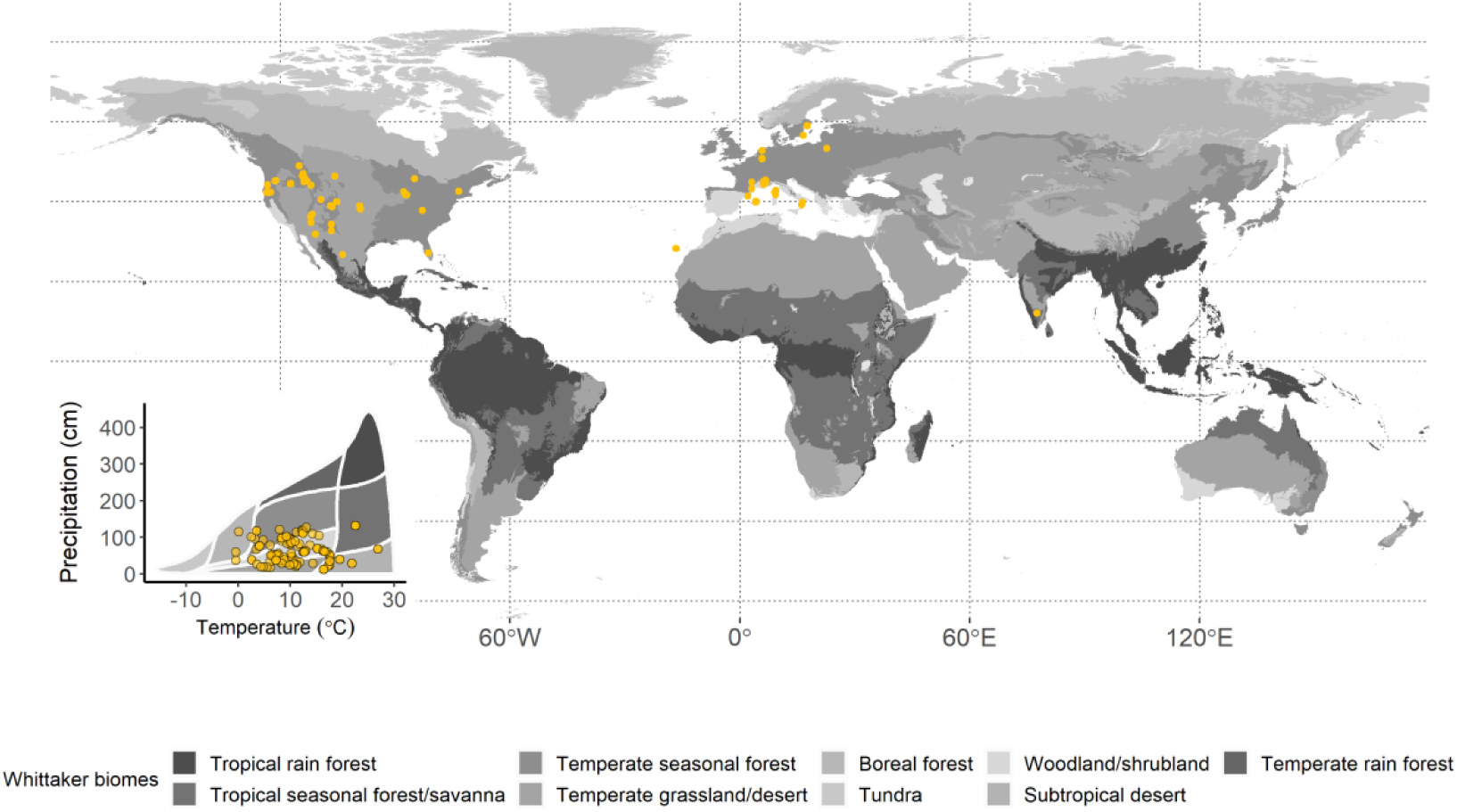
Geographic and locations and position in a Whittaker biome plot (insert) of the plant demographic studies used in this manuscript. The background grey scale represents Whittaker (1970) terrestrial biomes.

## Appendix S1. Creation of density-independent matrix population models from the data by Chu et al. (2016)

We used data published by Chu *et al*. (2016) to produce density-independent MPMs. To produce these MPMs, we fit statistical models as similar as possible to those used by Chu *et al*. (2016). The main difference is that our models did not include density-dependent terms. Below, we provide a brief explanation of the dataset, model fitting, and matrix projection construction.

The data from Chu *et al*. (2016) provide information on the survival, growth, and recruitment of 12 species across five study sites. These study sites are located in Arizona, Idaho, Kansas, Montana, and New Mexico. Each site contains a set of one-square-meter plots subdivided in groups of plots that are spatially adjacent. Each plot was censused annually using a pantograph (Hill 1920) that records the shape and position of each individual, classified as either adult or recruit. From the information on the shape of individuals, it is straightforward to infer the area of individuals; specifically, basal area for the grasses, and canopy area for the shrubs. These charts were digitized as shapefiles, and made public in a series of separate data articles (Adler *et al*. 2007; Zachmann *et al*. 2010; Anderson *et al*. 2011, 2012). These data are most commonly used to estimate the survival and growth of individuals, and the plot-level recruitment of seedlings (e.g. Adler *et al*. 2010; Dalgleish *et al*. 2011; Tredennick *et al*. 2018). The censuses extend from a minimum of 14 years at the Montana site, to a maximum of 35 years at the Kansas site.

We used these data to parameterize density-independent MPMs by fitting generalized linear mixed models to the survival, growth, and recruitment data. These models follow closely those fit by Dalgleish *et al*. (2011) on the subset of data from Idaho (Zachmann *et al*. 2010). In the individual models on survival and growth, we used the natural logarithm of basal or canopy area as our measure of individual size (henceforth “*log(size)*”). These models included random intercepts of year and group, and a random year slope of *log(size)*. As reported elsewhere (Tredennick *et al*. 2018), in these data, model selection supports model fits with both a random intercept, and a random slope of *log(size)*. We fit the survival model assuming a Bernoulli distributed response. Our survival model thus had the form

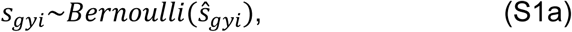

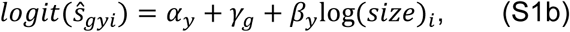

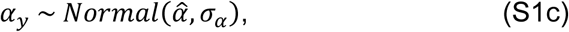

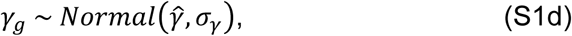

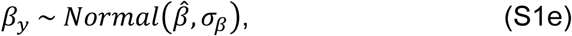

where *g* refers to group, *y* refers to year, *i* refers to individual observations, *α*_*y*_ is the random effect of year, *γ*_*g*_ is the random effect of group, *β*_*y*_ is the random year effect of size, and *ŝ*_*gyi*_ is predicted survival for each data point to time *y*. Here, note that *log(size)*_*I*_ refers to time *y-1*, so that *log(size)* of individual *i* in year *y-1* is used to predict the average survival (*ŝ*_*gyi*_) of individual *i* to year *y*.

The growth model had an identical structure, but it was a linear mixed effect model with normal response, so that:

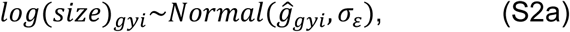

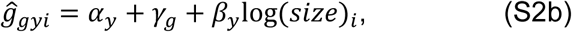

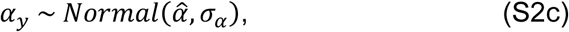

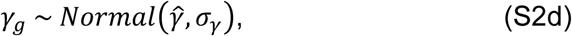

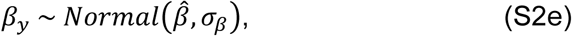

where *ĝ*_*gyi*_ is the predicted *log(size)* at time *y, log(size)*_*i*_ refers to time *y-1, σ*_*ε*_ is the residual standard deviation of the normally distributed variable, and all other symbols reflect the same model structure as found in the survival model (Eq. S2). In these data, the residual variance is not constant but depends on size. We modeled this size dependence on the residuals, defined as *r*_*gyi*_ = *log*(*size*)_*gyi*_ − *ĝ*_*gyi*_, with the nonlinear function

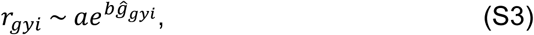

which models the residual of observation of site *g*, year *y*, and individual *i* as a nonlinear function of the corresponding predicted size, *ĝ*_*gyi*_, and parameters *a* and *b*. We fit the survival models in Eq. S1 using function glmer, the growth models in Eq. S2 using function lmer, both from the R package lme4 (Bates *et al*. 2015), and we fit Eq. S3 using the R function nls (R Core Team 2019).

Because recruitment data were available at the plot level, we used a different model structure. We used two predictors of plot-level recruitment: the plot-level and group-level cover of parents, following the model in Dalgleish *et al*. (2011). This model assumes that the number of recruits in year *y* and plot *i* reflects the cover of parents in the same plot *i* in year *y-1*, and dispersal from outside the plot, which should be proportional to the average cover of parents within the group of plots to which plot *i* belongs. Hence, we modeled recruitment of plot *i* in year *y* based on the cover of adults in the previous year, *cov*_*i*_, and the average cover of adults in the group of plots to which plot *i* belongs, *cov*_*g*_. We also modeled recruitment based on random effects of year and population. We modeled recruitment as a negative binomial process with a log link,

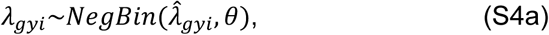

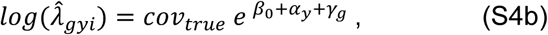

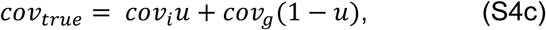

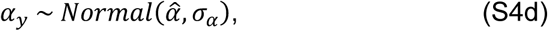

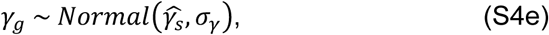

where θ is the overdispersion parameter, 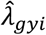 is the average predicted number of recruits in plot *i*, group *g*, and year *y, u* is a parameter bounded between 0 and 1 that quantifies the contribution of the cover from each individual plot *i*, and the average cover of the group *g*, to the number of recruits of the following year, *α*_*y*_ is a random effect of year *y, γ*_*g*_ is a random effect of group *g*, and *β*_0_ is the fixed effect of true cover.

We fit this model in a Bayesian framework, using Gibbs sampling via the R package R2jags (Yu-Sung & Yajima 2015). We ran three chains for 25,000 iterations, with 5,000 iterations of burn in. We assumed parameters had converged if the Gelman-Rubin metric 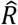 was below 1.1. Finally, we checked our models performing posterior predictive checks (Gelman *et al*. 2013).

We used these models to parameterize MPMs for each group and year. We created 10×10 MPMs, as suggested by Salguero-Gómez & Plotkin (2010). Our approach was similar to the construction of an IPM, in that we populated our MPMs using the parameters of the mixed effect models. However, IPMs usually model the size distribution of seedlings using a normal distribution. We did not use this method, because it would unintentionally evict most seedlings (Williams *et al*. 2012). Instead, we placed all recruits in the smallest size class.

## Appendix S2. The relationship between WAI and the effect of precipitation assuming the per-mm sensitivity of plant population growth does not change with WAI

We use a simulation to show that if the per-mm sensitivity of population growth rate to precipitation anomalies were the same across our 165 populations, the relationship shown in Figure 1A would be positive rather than negative. We focus on Figure 1A, because the artifact will be stronger when testing *H*_*2*_, and because *H*_*2*_ was partially supported by data. In *H*_*2*_, we posited that the *λ* of plants in water-limited biomes would be more responsive to precipitation anomalies. Therefore, we expected that the response of plant population growth rate to standardized precipitation anomalies should decrease with water availability index (WAI). This hypothesis implicitly assumes that the per-mm sensitivity to precipitation, or, in other words, the sensitivity to *absolute* precipitation anomalies, increases with site aridity (Huxman *et al*. 2004). However, if the sensitivity to absolute precipitation anomalies were the same across the WAI gradient, the meta-regression Figure 1A would show a positive, rather than negative correlation. This would occur because WAI is strongly correlated with the standard deviation of *absolute* precipitation anomalies (Fig. S2A). As a result, at our most and least arid sites, a *standardized* precipitation anomaly of 1 corresponds to a standard deviation of 47mm and 181mm, respectively. Hence, if the per-mm sensitivity to absolute precipitation anomalies were the same across this WAI gradient, we would find a positive relationship, rather than the negative relationship we found in Fig. 1A of the main text.

To demonstrate this statistical artifact, we performed a simulation study using parameters consistent with the data we used in our study. Specifically, we simulated 100 time series of 20 values, each corresponding to a value on a gradient of 100 WAI values. These 100 datasets simulated the process

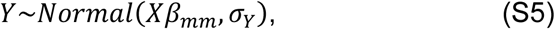

where *Y* represented *log(λ)* in our analyses, *X* were the absolute precipitation anomalies, *β*_*mm*_ the average effect size of absolute precipitation anomalies, and *σ*_*Y*_ was the residual variance. We set *β*_*mm*_ and *σ*_*Y*_ using the coefficients estimated via Eq. 1 in the main text. However, the *β* estimated by Eq. 1, which was equal to 0.032, estimated the effect of *standardized* rather than *absolute* precipitation anomalies on *log(λ)*. To simulate the process in Eq. S5, we therefore converted *β* to a per-mm effect of precipitation of *β*_*mm*_=0.0003mm. We obtained this number assuming *β* referred to our median WAI of - 309mm, and therefore to a standard deviation of 115mm (Fig. S2A) and a per-mm effect of absolute precipitation anomalies of *β*_*mm*_=0.032/115mm=0.0003mm. We set *σ*_*Y*_ to 0.16, by calculating the average *ε* value estimated fitting Eq. 2 and 3 of the main text across our 165 populations. Finally, we simulated the *X* values corresponding to each one of the 100 points across the WAI as

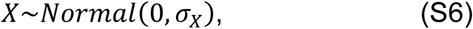

where *σ*_*X*_ was the expected absolute (in mm) standard deviation in annual precipitation at each one of our 100 WAI values (the line in Fig. S2A). We used these X values to simulate the Y values according to Eq. S5. For each one of these 100 datasets, we estimated the linear model *Y* = *β*_*r*_*A* + *ε*, where *A* is the *standardized* anomaly given by *A = E(X)/σ(X)*.

Plotting the *β*_*r*_ coefficients versus the corresponding WAI-values provides a positive, rather than negative relationship (Fig. S3). This positive trend in sensitivities results because the *standardized* anomalies, *A*, refer to larger and larger absolute anomalies *X* generating the data.

**Figure S2.**
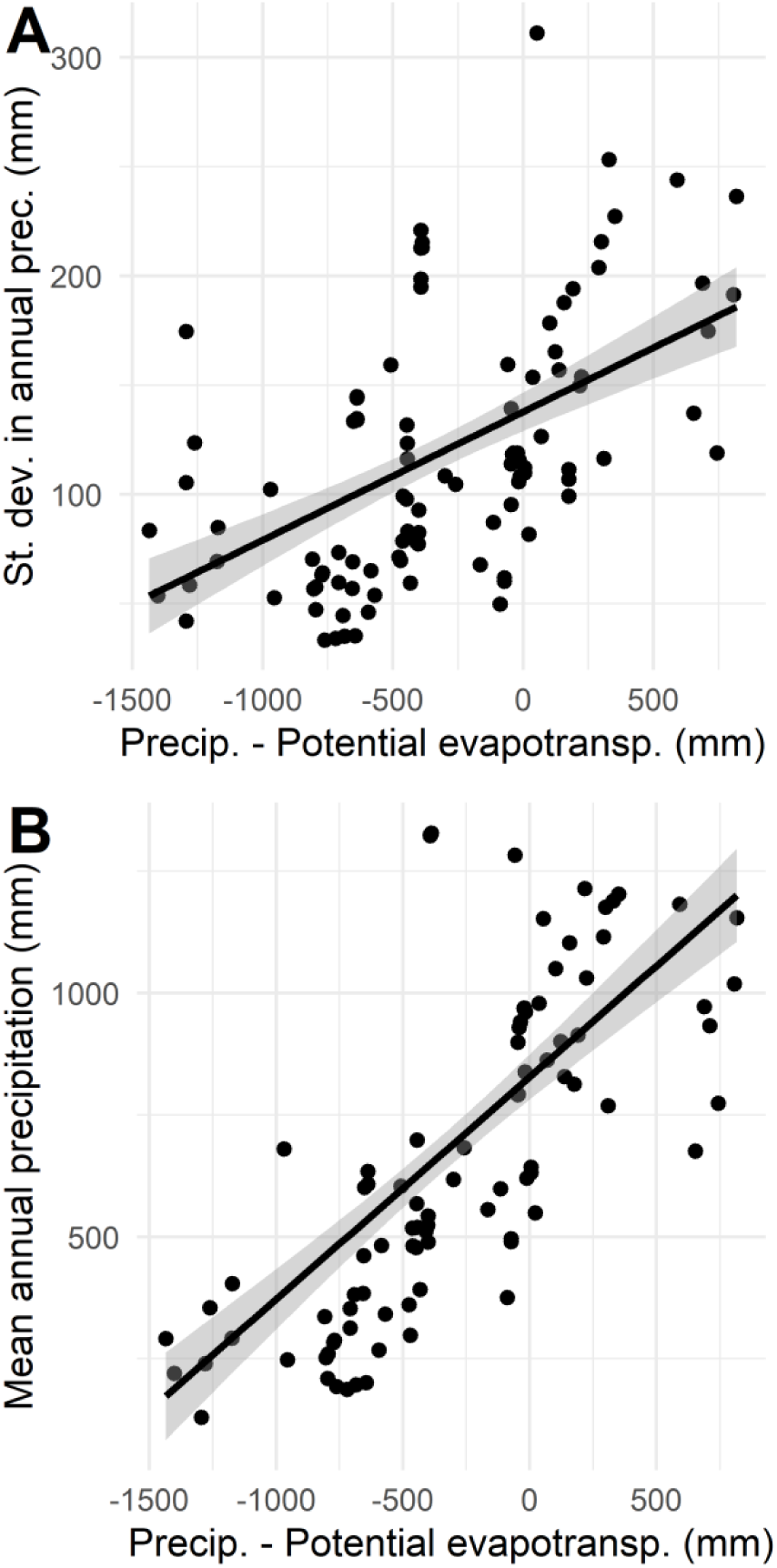
Relationship between water availability index (WAI), the standard deviation of precipitation, and mean annual precipitation. The standard deviation refers to a 40-year period of annual precipitation data. Each point refers to one of the 165 plant populations included in our study.

**Figure S3.**
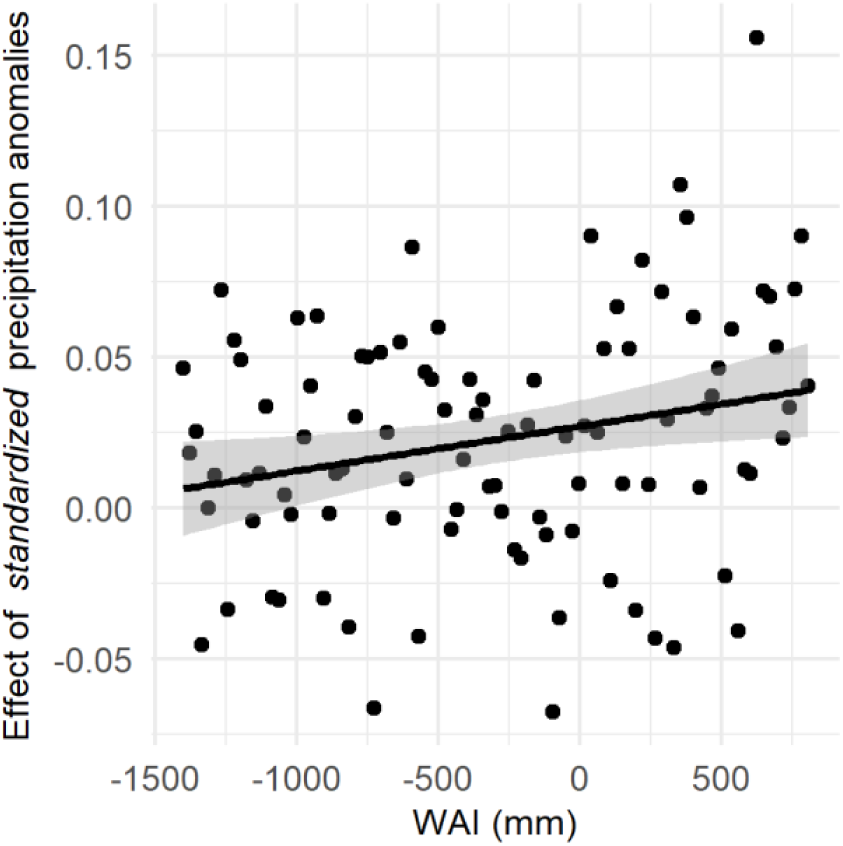
Meta-regression showing the effect sizes recovered from a simulation assuming a constant per-mm effect of precipitation across WAI-values. The x-axis shows 100 equally spaced WAI-values. The y-axis shows effect sizes akin to the ones we extracted in the meta-regression represented in Figure 1A of the main text.

## Appendix S3. Methods for computing generation time and the degree of iteroparity

We calculated generation time (*T*) and the degree of iteroparity (S) using methods described in Salguero-Gómez *et al*. (2016). We calculated generation time (*T*) by applying the age-specific survivorship and fertility curves

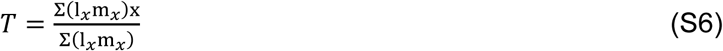

where *l*_*x*_ and *m*_*x*_ are, respectively, survivorship and fertility at age *x*. These two age-specific functions were obtained using age-from-stage methods defined by Cochran & Ellner (1992) and Caswell (2001). The vectors *l*_*x*_ and *m*_*x*_ were trimmed at the quasi-stable stage of 95% convergence to the stable stage distribution to avoid potential spurious plateaus of mortality and fertility (Horvitz & Tuljapurkar 2008) as described in Jones *et al*. (2014).

We calculated the degree of iteroparity (*S*) following the formula for Demetrius’s entropy (1978). This measure of entropy relies again on the vectors *l*_*x*_ and *m*_*x*_ described above, and on *λ*, the asymptotic population growth rate, calculated as the dominant eigenvalue of the transition matrix ***A*** (Caswell 2001):

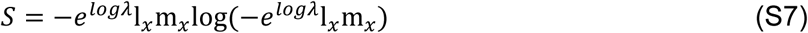

Values of *S* close to 0 correspond to strictly semelparous species, whereas greater values of *S* describe greater degrees of iteroparity, and thus longer reproductive windows.

## Appendix S4. Calculating the proportion of the climate covered by each study

To calculate the proportion of climate variation observed during each climate driver study, we represented the climate at each population site using two-dimensional convex hulls. We defined “climate variation” as the two-dimensional convex hull delimited by the combination of annual total precipitation, and annual mean temperature observed during a reference time period. Therefore, the larger the range of precipitation and temperature observed during a time period, the larger the area of the convex hull. For each population, we calculated the convex hulls referred to the period of demographic census, and to the 40-year period representative of climate at each site. This 40-year period ends at the last demographic census of each study. By definition, the convex hull referring to the study period is a subset of the convex hull referring to the 40-year period representative of climate (Figure S4). We calculated the area of these convex hulls using functions *chull* in base R, and *Polygon* and *SpatialPolygons* from package sp (Bivand *et al*. 2013).

**Figure S4.**
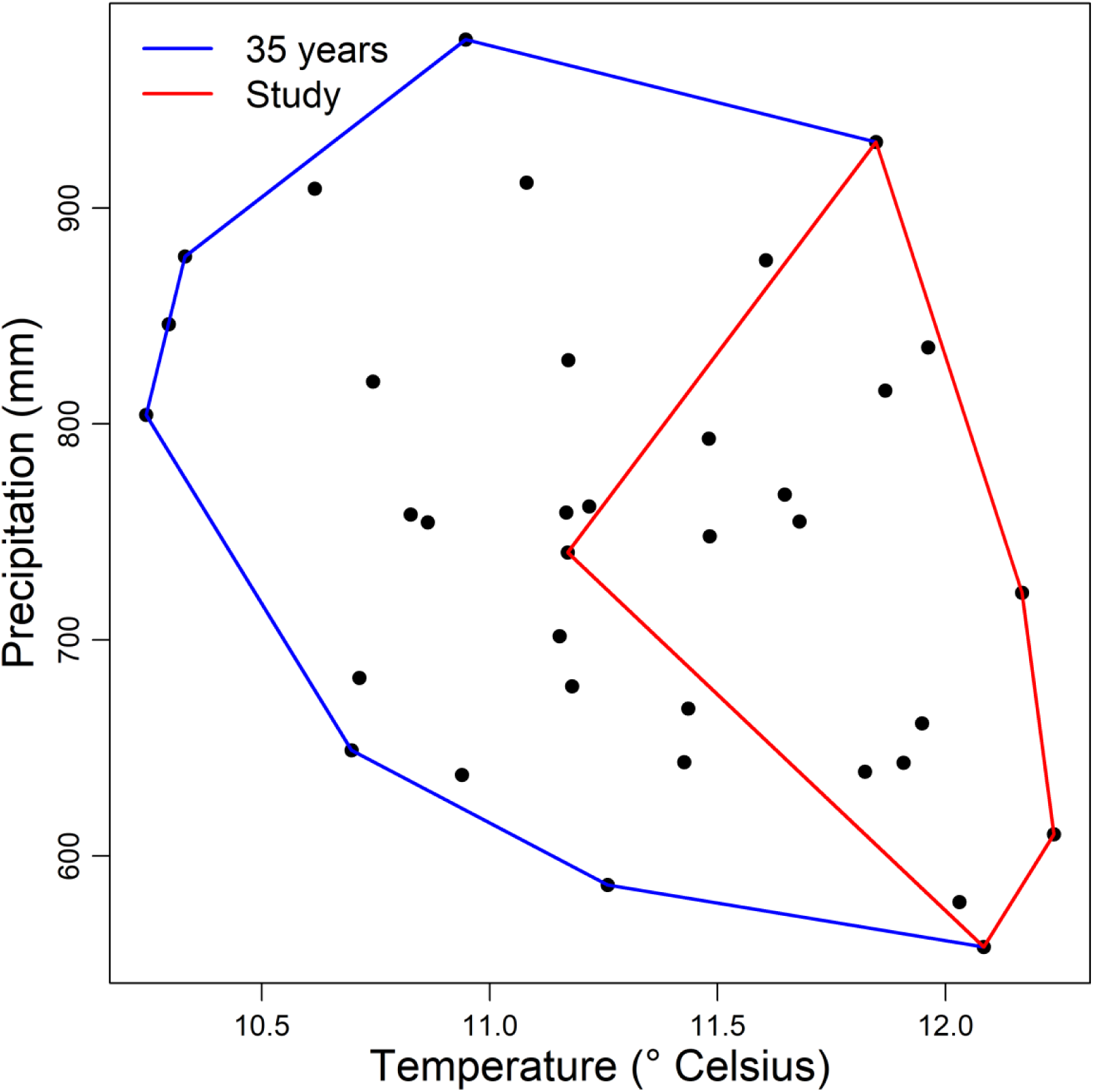
Example of convex hulls showing the range of climate observed during a demographic study (red) and during the 40-year period representative of climate (blue). This example is referred to the study carried out on *Brassica insularis* by Noël *et al*. (2010).

